# An Evaluation of the *In Vitro* Roles and Mechanisms of Silibinin in Reducing Pyrazinamide- and Isoniazid-Induced Hepatocellular Damage

**DOI:** 10.1101/815241

**Authors:** Zhang-He Goh, Jie Kai Tee, Han Kiat Ho

**Author notes:** Corresponding author at: Department of Pharmacy, Faculty of Science, National University of Singapore, 18 Science Drive 4, Singapore 117543, E-mail address (H.K. Ho).

## Abstract

Tuberculosis remains a significant infectious lung disease that affects millions of patients worldwide. Despite numerous existing drug regimens for tuberculosis, Drug-Induced Liver Injury is a major challenge that limits the effectiveness of these therapeutics. Two drugs that form the backbone of the commonly administered quadruple antitubercular regimen, i.e. pyrazinamide (PZA) and isoniazid (INH), are associated with such hepatotoxicity. The problem is compounded by the lack of safe and effective alternatives to the antitubercular regimen. Consequently, current research largely focuses on exploiting the hepatoprotective effect of nutraceutical compounds as complementary therapy. Silibinin, a herbal product widely believed to protect against various liver diseases, potentially provides a useful solution given its hepatoprotective mechanisms. In our study, we identified silibinin’s role in mitigating PZA- and INH-induced hepatotoxicity and elucidated a deeper mechanistic understanding of silibinin’s hepatoprotective ability. 25 μM silibinin preserved the viability of human foetal hepatocyte line LO2 when co-administered with 80 mM INH and decreased apoptosis induced by a combination of 40 mM INH and 10 mM PZA by reducing oxidative damage to mitochondria, proteins, and lipids. Taken together, this proof-of-concept forms the rational basis for the further investigation of silibinin’s hepatoprotective effect in subsequent preclinical studies and clinical trials.

**Graphical Abstract:** 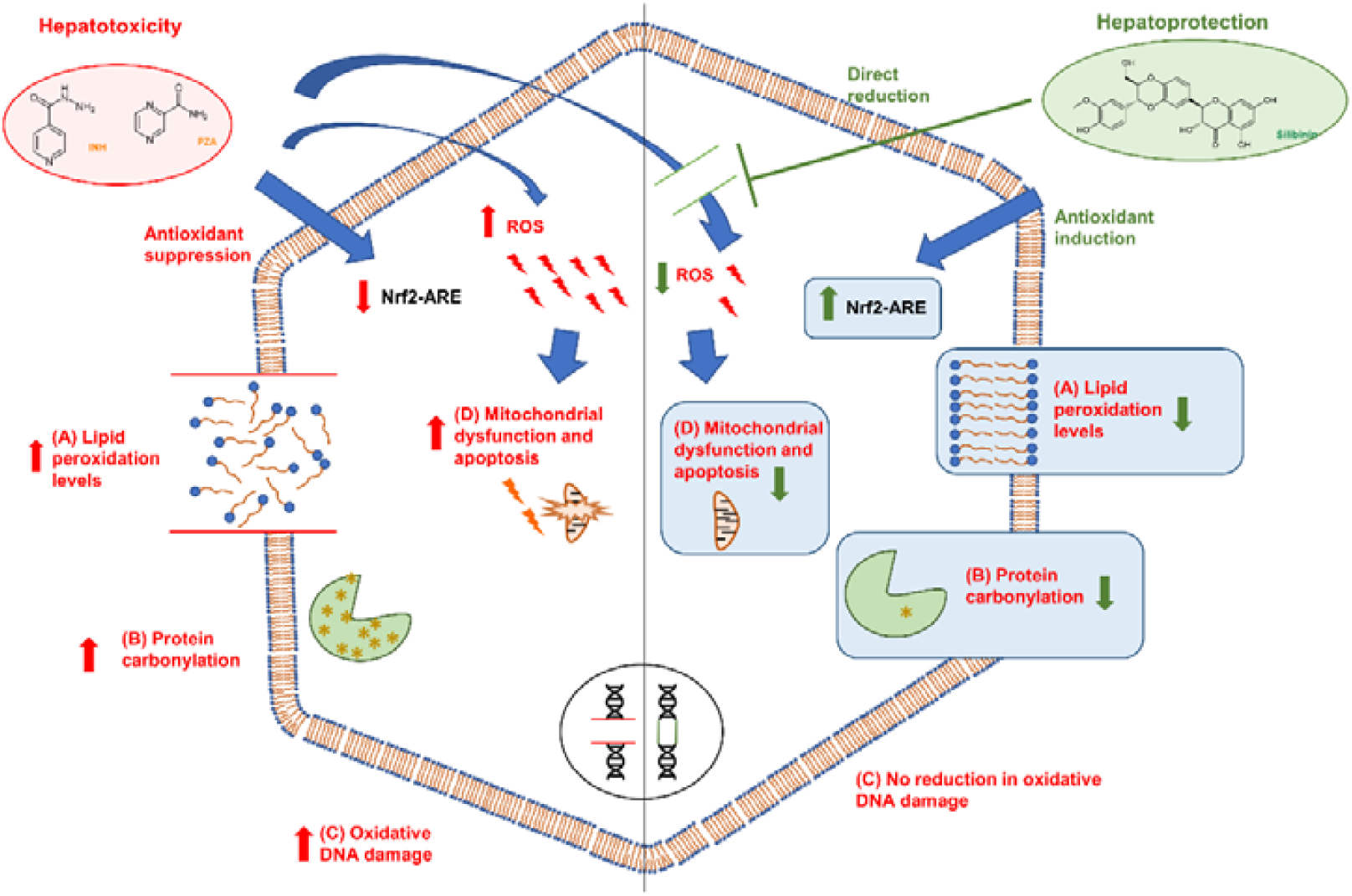

## 1. Introduction

Tuberculosis is an infectious lung disease caused by *Mycobacterium tuberculosis*, with one in three people affected globally (Global Tuberculosis Report, 2018). While new and current drug regimens have been tailored to shorten treatment duration and increase therapeutic efficacy (Kwon et al., 2014), Drug-Induced Liver Injury (DILI) caused by anti-tubercular therapy (ATT) still warrants the most concern. ATT has been reported to cause severe hepatotoxicity (Abbara et al., 2017; Cano-Paniagua et al., 2019), which leads to discontinuation of therapy in 11% of patients (Ramappa and Aithal, 2013). Hence, the DILI arising from ATT is a pressing problem that needs to be addressed in tuberculosis.

The most prevalent ATT, known as the PIER regimen, involves the use of four drugs in combination: pyrazinamide (PZA), isoniazid (INH), ethambutol (EMB), and rifampicin (RMP). Among them, PZA and INH are the most commonly implicated in DILI. PZA increases the risk of DILI by 3.5 times (Chang et al., 2008). In HepG2, PZA-induced hepatotoxicity has been reported to invoke damage to cellular and mitochondrial membranes, leading to increased apoptotic activity *in vitro* (Singh et al., 2012). Similarly, INH-induced hepatotoxicity occurs in approximately 25% of patients (Wang et al., 2016a), and severe INH-induced hepatotoxicity occurs in 1 per 1000 patients (Harrington et al., 2010). At the same time, rat studies have also been conducted on PZA, INH, and RMP, as well as various combinations of these drugs. Administration of these combinations have led to increased membrane lipid peroxidation levels (Eminzade et al., 2008), increased serum levels of liver enzymes (Srivastava et al., 2008; Tasduq et al., 2005), and reduced antioxidant protein levels (Tasduq et al., 2005; Victorrajmohan et al., 2005). Notably, the search for strategies to reduce PZA- and INH-induced hepatotoxicity is further underscored by the first-line status of the PIER regimen in ATT with few safer and equally efficacious alternatives (Jiménez-Arellanes et al., 2016; Zumla et al., 2013).

The two main mechanisms through which PZA and INH injure hepatocytes are both associated with oxidative stress (Betteridge, 2000). First, PZA and INH can be converted to reactive metabolites by drug metabolizing enzymes. PZA is oxidized to 5-hydroxypyrazinamide via xanthine oxidase; both PZA and 5-hydroxypyrazinamide are further bioactivated by the same enzyme to the toxic metabolites, pyrazinoic acid and 5-hydroxypyrazinoic acid (Pitrè et al., 1981; Shih et al., 2013). 5-hydroxypyrazinoic acid has been hypothesised to be the more toxic of the two active metabolites (Shih et al., 2013). In contrast, INH can be activated to toxic metabolites via N-acetyltransferase and amidases (Liu et al., 2017; Tafazoli et al., 2008; Wang et al., 2016a), as well as CYP2E1 (Liu et al., 2017; Wang et al., 2016a), the latter of which INH also induces (Hassan et al., 2018). These metabolites increase the levels of intracellular ROS (Ahadpour et al., 2016; Sharma et al., 2018; Wang et al., 2016a), thus damaging vital cellular targets. The subcellular consequences include: DNA fragmentation (Sharma et al., 2018), lipid peroxidation (Cao et al., 2018; Rawat et al., 2018; Sharma et al., 2018), and protein carbonylation (Rawat et al., 2018; Tafazoli et al., 2008). Therefore, given that the multifactorial nature of DILI arising from ATT includes interindividual variability in metabolic processes, patients may exhibit features of idiosyncratic toxicity.

Second, PZA and INH suppress the Nuclear factor (erythroid-derived 2)-like 2 (Nrf2)-antioxidant response element (ARE) pathway that protects cells from oxidative damage (Chen et al., 2013; Peng et al., 2016). Consequently, both PZA (Peng et al., 2016) and INH (Chen et al., 2013) increase cellular susceptibility to oxidative stress by decreasing the expression of downstream antioxidant proteins; the antioxidant proteins affected include Glutamate-cysteine ligase catalytic subunit (Gclc), NAD(P)H quinone dehydrogenase 1 (NQO1), Heme oxygenase-1 (HO-1), and Sulfiredoxin 1 (Srxn1). Indeed, RMP, INH, and PZA have been shown to potentiate the hepatotoxic effects of one another in *in vitro* assays involving HepG2 (Singh et al., 2011), though the underlying mechanisms of hepatotoxicity caused by these antitubercular drugs remain poorly understood to date. Seen in totality, the broad strokes illustrated by these studies denotes the need for hepatoprotective strategies to counter the increase in oxidative stress induced by these drugs.

The most well-researched strategy to reduce PIER-induced hepatotoxicity involves the use of antioxidant nutraceuticals to reduce oxidative stress and liver inflammation (Li et al., 2015). Among the nutraceuticals that have been explored for their hepatoprotective potential, silibinin is the gold standard (Surai, 2015). Silibinin is a herbal product derived from milk thistle that has been postulated to protect against liver injury caused by various chemotherapeutic (Patel et al., 2010) and toxic agents (Fanoudi et al., 2018; Jiménez-Arellanes et al., 2016; Ma et al., 2016). Two factors contribute to silibinin’s popularity over other chemical drugs in liver disease: it has low toxicity (Flora et al., 1998) and exhibits a broad spectrum of hepatoprotective mechanisms (Polachi et al., 2016; Rodriguez-Garcia et al., 2017; Vargas-Mendoza et al., 2014). Many of silibinin’s mechanisms of action can be attributed to its antioxidant, anti-inflammatory, immunomodulatory, and antifibrotic actions (Abenavoli et al., 2018). Silibinin’s anti-inflammatory and immunomodulatory effects are manifested through silibinin’s actions on pathways involving tumor necrosis factor-alpha (TNF-α) (Abdel-Moneim et al., 2015) and NF-κB (Kim et al., 2015), as well as its modulation of lipopolysaccharide-induced NO production (Kim et al., 2015) and NLRP3 inflammasome activation (Zhang et al., 2018). At the same time, silibinin’s antioxidant effect has been attributed to its ability to inhibit ROS-producing enzymes, directly scavenge free radicals, prevent the absorption of ions by the intestine through chelation, and promote the expression of protective molecules and enzymes that mitigate oxidative stress (Abenavoli et al., 2018; Surai, 2015). Therefore, silibinin may be uniquely placed to mitigate the principal mechanisms implicated in oxidative stress responsible for PIER-induced hepatotoxicity.

Unfortunately, despite the extensive and rigorous research on the basis for silibinin’s hepatoprotective effect in recent years (Ramappa and Aithal, 2013; Roubalová et al., 2017; Surai, 2015), the *in vivo* and *in vitro* biological markers which silibinin modulates have not been conclusively linked to the reduction of DILI (Mann et al., 2017). Furthermore, the exact biochemical mediators behind silibinin’s hepatoprotective effect have also not been identified (Raghu and Karthikeyan, 2016). Silibinin’s hepatoprotective nature remains nebulous to date, making it especially challenging to clarify and optimise silibinin’s role in mitigating PIER-induced hepatotoxicity. Consequently, silibinin’s reduction of PIER-induced hepatotoxicity has neither been definitively proven nor characterised (Luangchosiri et al., 2015; Marjani et al., 2016). Therefore, a deeper mechanistic understanding of silibinin’s hepatoprotective ability must be elucidated before silibinin can be widely used as an adjuvant to ameliorate PIER-induced hepatotoxicity.

In this study, we sought to investigate the role of silibinin in mitigating PZA- and INH-induced hepatotoxicity. We hypothesised that silibinin reduces PZA- and INH-induced hepatotoxicity through its anti-oxidative mechanisms. Indeed, our results showed that silibinin preserved cell viability when co-administered with INH. We also determined that the co-administration of silibinin with a combination of INH and PZA (I/P) led to a reduction in oxidative damage to intracellular targets and apoptotic activity. Together, these findings supported our hypothesis that silibinin reduces PZA- and INH-induced hepatotoxicity through its modulation of oxidative stress.

## 2. Materials and Methods

### 2.1 Cell culture and reagents

LO2 is a human foetal hepatocyte cell line that has been previously characterised (Hu et al., 2013). TAMH was a kind gift from the late Prof. Nelson Fausto (University of Washington); the isolation of TAMH was previously described (Wu et al., 1994). LO2 was cultured in Dulbecco’s minimum essential medium (DMEM) (Sigma Aldrich, United States) containing 10% v/v foetal bovine serum (FBS). TAMH was cultured in DMEM-F12 (Sigma Aldrich, United States). Cells were incubated at 37 °C in a humidified incubator with 5% CO_2_. Stocks of 100 mM silibinin (Sigma Aldrich, United States), 10 mM sulphoraphane (SU) (Sigma Aldrich, United States), 50 mM trans-cinnamaldehyde (CA) (Sigma Aldrich, United States), and 5 M tert-butyl hydroperoxide (TBHP) (Sigma Aldrich, United States) were prepared in dimethyl sulfoxide (DMSO) (Sigma Aldrich, United States). Stocks were diluted with culture medium into different concentrations, ensuring that the final concentration of DMSO never exceeded 0.1% v/v.

### 2.2 Cell viability assay

10,000 cells per well were seeded in a clear 96-well plate overnight and treated accordingly. At each timepoint, 0.5 mg/mL 3-(4,5-dimethylthiazol-2-yl)-2,5-diphenyltetrazolium bromide (MTT) solution in fresh media was added to the cells and incubated for 3 h at 37 °C. Thereafter, the MTT solution was removed, the formazan crystals formed dissolved in 200 μL DMSO, and the absorbance measured at 570 nm using Hidex sense microplate reader (Hidex, Finland).

### 2.3 Direct ROS quantitation

10,000 cells per well were seeded in a black 96-well plate overnight and treated accordingly. At each timepoint, cells were washed with phosphate-buffered saline (PBS) and incubated for 30 min with 10 **µ**M 6-chloromethyl-2’,7’-dichlorodihydrofluorescein diacetate (DCFDA) and 1mg/L Hoechst 33342 dye (Sigma Aldrich, United States) diluted in media. 100 **µ**L of PBS was added and fluorescence measured at **λ**_**ex**_**/λ**_**em**_ **=** 350/461 nm (Hoescht) and 485/535 nm (DCFDA) respectively using Hidex sense microplate reader (Hidex, Finland).

### 2.4 Lipid peroxidation quantitation

20,000 cells/cm^2^ were seeded onto a 100 mm dish overnight and the respective treatment media was added. Thiobarbituric acid reactive substances (TBARS) were quantified using the TBARS assay kit (Cayman Chemical, United States) following the manufacturer’s instructions. Briefly, both the adhered live cells and floating dead cells were harvested, resuspended in 150 µL PBS, and sonicated for 5 min. 100 µL of resuspended pellet and 100 µL SDS solution were added to test tubes and mixed with 4 mL of Colour Reagent solution, which constituted 530 mg of 2-thiobarbituric acid dissolved in 50 mL of a 1:1 mixture of 20% v/v acetic acid and 10% w/v sodium hydroxide. The tubes were boiled for 1 h, immersed in ice for 10 min to quench further reaction, centrifuged at 2000 *g* for 10 min at 4 °C, warmed to room temperature, and added to a 96-well black plate. Fluorescence intensity was measured at **λ**_**ex**_**/λ**_**em**_ **=** 520/560 nm using Hidex sense microplate reader (Hidex, Finland). Lipid peroxidation levels were normalised against total protein quantitated using the Pierce BCA Protein Assay Kit (Thermo Fisher Scientific, United States).

### 2.5 Protein carbonylation quantitation

20,000 cells/cm^2^ were seeded onto a 100 mm dish overnight and treated accordingly. Both live and dead cells were harvested and resuspended in 150 **µ**L MilliQ Grade I water. 100 **µ**L of this suspension was added to 500 **µ**L 10% v/v trichloroacetic acid (TCA) solution and centrifuged at 13,000 rpm for 2 min. The cell pellet was collected and incubated with 100 **µ**L 0.02% w/v 2,4-dinitrophenylhydrazine (2,4-DNPH) hydrochloride (Tokyo Chemical Industries, Japan) for 1 h with constant vortexing. 50 **µ**L 100% v/v TCA was added and the suspension was centrifuged at 13,000 rpm for 5 min. The cell pellet was washed with cold acetone and dissolved in 200 **µ**L 6M guanidine HCl (Sigma Aldrich, United States). The absorbance of the solution was measured at 375 nm in Hidex sense microplate reader (Hidex, Finland) to determine carbonyl levels, which were normalised against total protein quantitated using the BCA assay.

### 2.6 Comet assay

400,000 cells per well were seeded on a 12-well plate overnight, treated accordingly, harvested and resuspended in 100 **µ**L PBS. 20 µL of this suspension was mixed with low-melting agarose (Trevigen, United States), spread evenly over CometSlides (Trevigen, United States), left to congeal, then kept in lysis solution (Trevigen, United States) at 4 °C overnight. Thereafter, the slides were immersed in unwinding solution for 30 min at room temperature before gel electrophoresis was run for 25 min. The slides were washed, dried at 37 °C overnight, and stained with SYBRGold (Qiagen, Germany). Fluorescence images were taken with Olympus Fluoview FV1000 confocal microscope (Olympus, Japan) and analysed using OpenComet (Gyori et al., 2014). The Tail Moment and the Olive Moment were calculated as follows:

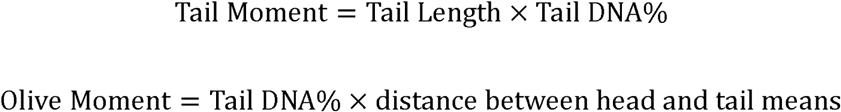

### 2.7 Mitochondrial membrane potential measurement

20,000 cells/cm^2^ were seeded onto a 60 mm dish overnight and treated accordingly. They were then incubated with Tetramethylrhodamine-methyl-ester (TMRM) (Thermo Fisher Scientific, United States) for 30 min, harvested, centrifuged, and reconstituted in 400 **µ**L PBS. The percentage of cells within a defined range of fluorescence intensity was determined with Beckman Coulter CyAn ADP flow cytometer (Beckman Coulter, United States).

### 2.8 Apoptosis detection

20,000 cells/cm^2^ were seeded onto a 60 mm dish overnight and treated accordingly. Cell lysates were extracted with radioimmunoprecipitation assay (RIPA) buffer containing 0.1% w/v SDS, 1% w/v NP-40 and 0.5% w/v sodium deoxycholate in PBS. 300 µL of protease assay buffer (2 mM DTT, 10% v/v glycerol, 20 mM HEPES, 20 mM Ac-DEVD-AMC Caspase-3 Fluorogenic Substrate (BD Pharmingen, United States)) was added and the samples were incubated at 37 °C in the dark for 1 h. 100 µL of each sample was added to a 96-well black plate and the fluorescence intensity was measured at **λ**_**ex**_**/λ**_**em**_ **=** 390/444 nm using Hidex sense microplate reader (Hidex, Finland).

### 2.9 Western Blot

Cell lysates were extracted using RIPA buffer and protein concentrations were normalised using the BCA assay. Proteins were mixed with loading dye, boiled at 100°C for 5 min, separated on SDS-PAGE using 12% v/v polyacrylamide gels (Bio-Rad Laboratories, United States), then transferred onto polyvinylidene difluoride (PVDF) membrane (Thermo Fisher Scientific, United States) at 4°C with 100V for 2 h. Membranes were washed with Tris-buffered saline (1st Base, Singapore) containing 0.1% v/v Tween, blocked with 5% w/v bovine serum albumin (BSA), then incubated overnight at 4°C with the following primary antibodies in 2% w/v BSA: Rabbit anti-Gclc antibody (Abcam, United Kingdom; 1:1000); rabbit anti-NQO1 antibody (Cell Signalling, United States; 1:1000); rabbit anti-HO-1 antibody (Cell Signalling; United States; 1:1000); mouse anti-Srxn1 antibody (Santa Cruz, United States; 1:500); and mouse anti-**β**-actin antibody (Cell Signalling, United States; 1:10000). Bound antibodies were detected using horseradish peroxidase-conjugated secondary antibodies and visualized by chemiluminescence using Western Lightning Plus-ECL reagent (Perkin Elmer, United States). Band intensities were analysed using ImageJ (National Institutes of Health) and normalised using **β**-actin.

### 2.10 Statistical analysis

Statistical analysis was conducted using GraphPad Prism. Results were expressed as means ± S.E.M.. Differences in mean values were analysed by t-tests or one-way analysis of variance (ANOVA) with Tukey Honest Significant Difference test. A *p*-value of < 0.05 was considered statistically significant.

## 3. Results

### 3.1 Silibinin mitigated hepatotoxicity induced by INH when administered as a rescue adjuvant

As silibinin’s hepatoprotective role is often discussed in conjunction with its anticancer and antiproliferative properties (Bokemeyer et al., 1996; Elhag et al., 2015; Tiwari and Mishra, 2015), we first optimised silibinin’s treatment duration and established a suitable range of concentrations of silibinin that could be used safely without precipitating adverse effects. By testing the effects of various concentrations of silibinin on cell viability over 72 h, we determined silibinin’s maximum non-toxic concentration to be 50 **μ**M (**Fig. S1**). Consequently, subsequent experiments focused on testing silibinin’s hepatoprotective effect at the concentrations of 25 **μ**M and 50 **μ**M. Similarly, to optimise the concentration windows of INH and PZA, we determined their IC_50_s to be 73 and 60 mM respectively (**Fig. S2**).

Having established the optimal concentrations of silibinin, PZA, and INH to be used in our experiments, we then profiled silibinin’s orthogonal roles as a preventive, rescue, or recovery adjuvant in reducing PZA- and INH-associated hepatotoxicity. This approach was based on differential sequencing of the toxicant and silibinin exposure (**Fig. 1**). In our exploration of silibinin as a recovery adjuvant, we investigated silibinin’s ability to mitigate hepatocyte toxicity *in vitro* after hepatotoxic induction. We set up a pair of experiments, where on one hand, hepatocytes underwent a washout procedure to remove toxicant after 24 h, while on the other hand, the toxicant remained in the culture medium. By comparing silibinin’s *in vitro* hepatoprotective ability between this pair of experiments, we simulated silibinin’s potential ability to mitigate further liver injury in patients who either discontinue or stay on the hepatotoxic regimen.

**Figure 1.**
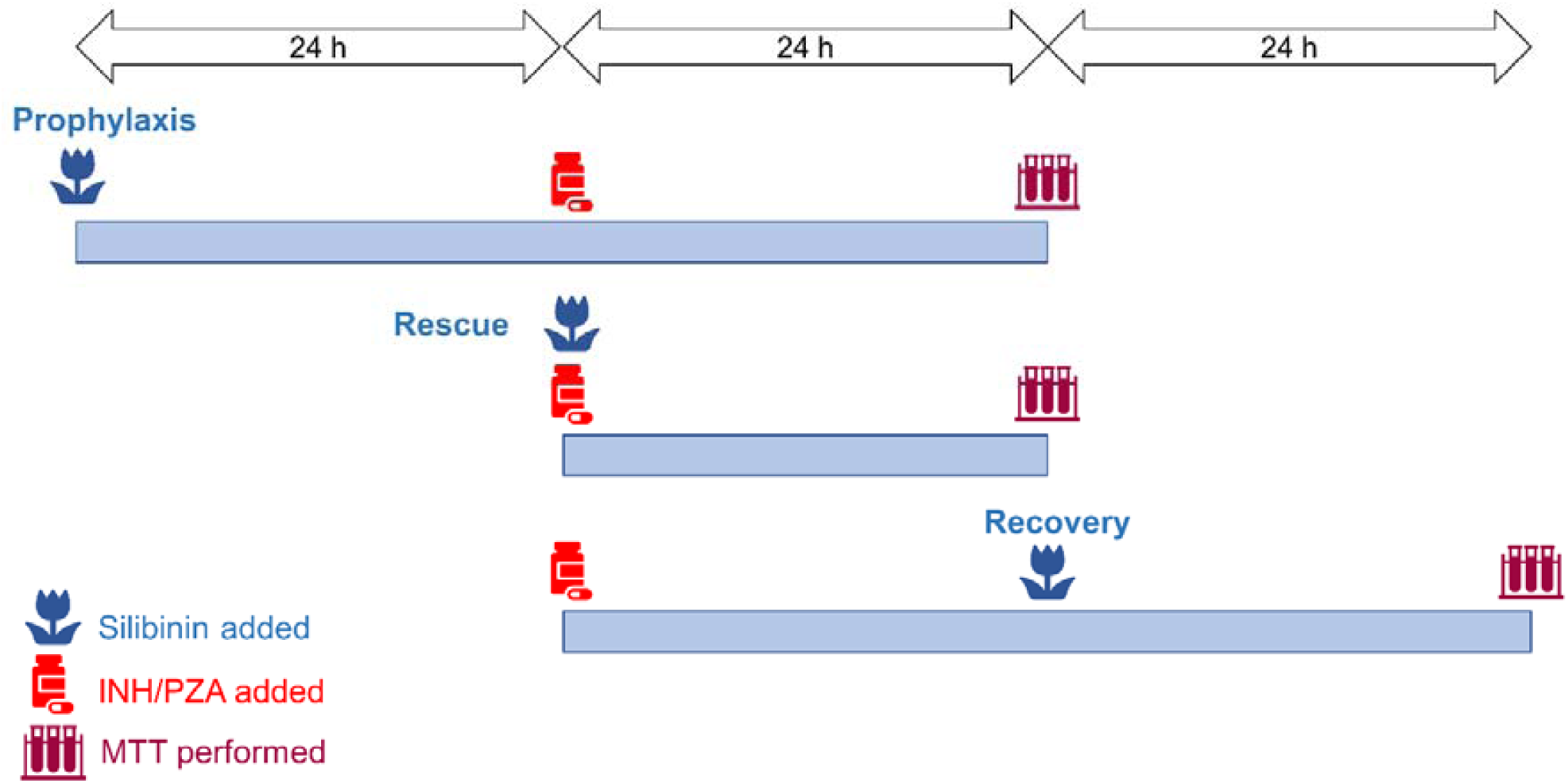
(color, 2-column). Treatment scheme involving silibinin’s role as a prophylactic, rescue, and recovery adjuvant. To simulate silibinin’s role as a preventive agent, silibinin was administered 24 h before the treatment with toxicants. To simulate silibinin’s role as rescue adjuvant, silibinin was co-administered with the toxicant regimen. To simulate silibinin’s role as a recovery adjuvant, silibinin was added 24 h after the toxicant regimen. The recovery experiments were further subdivided into two conditions: the first had a washout step, while the second did not have a washout step. In the simulation with washout, the toxicant regimen was replaced with silibinin alone and then treated for a further 24 h to investigate silibinin’s ability to aid patients in recovery after stopping the hepatotoxic regimen. In the simulation without washout, the toxicant regimen was replaced with a combination of silibinin and toxicant and treated for a further 24 h to investigate silibinin’s ability to mitigate further liver injury in patients who stay on the toxicant regimen.

Three major observations can be made about our experiments that serve to identify silibinin’s role in protecting against DILI. First, silibinin was effective in rescue (**Fig. 2A**), but not in prevention and recovery (**Fig. 2B-C**). The co-administration of 25 **μ**M silibinin with 80 mM INH moderately protected against INH-induced hepatotoxicity, raising the mean hepatocyte viability from 53% to 63% (**Fig. 2A**). Second, when hepatotoxicity was induced by a higher concentration of INH at 100 mM, the magnitude of silibinin’s hepatoprotective effect decreased slightly and silibinin’s optimal hepatoprotective concentration rose to 50 **μ**M (**Fig. 2A**). Inducing hepatotoxicity using a lower concentration of INH at 50 mM also appeared to negate silibinin’s hepatoprotective effect (**Fig. 2A**). Third, silibinin’s hepatoprotective effect was independent of PZA-induced hepatotoxicity *in vitro* (**Fig. 2A-C**), and silibinin’s protection against INH-induced hepatotoxicity lessened slightly when silibinin was administered together with a combination of 50 mM INH and 50 mM PZA (**Fig. 2A**). Everything considered, these results suggest that most of silibinin’s hepatoprotective effect may be the most apparent at moderate levels of INH-mediated toxicity.

**Figure 2.**
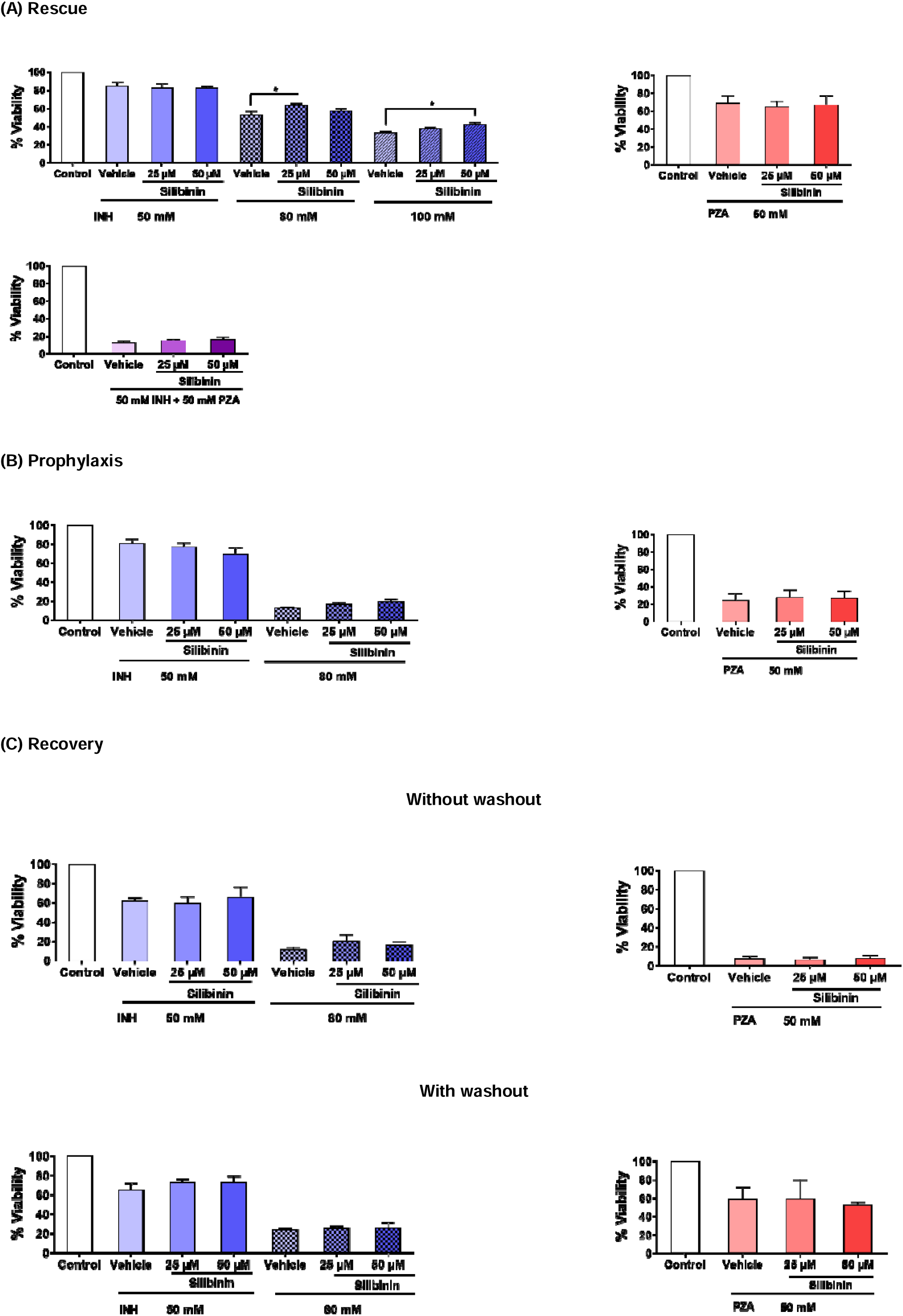
(color, 2-column). Silibinin mitigated INH-induced hepatotoxicity, but not PZA-induced hepatotoxicity. **(A)** Co-administration of silibinin at 25 µM reduced hepatotoxicity induced by 80 mM (one-way ANOVA, *p* = 0.010). Similarly, co-administration of silibinin at 50 µM reduced hepatotoxicity induced by 100 mM INH (one-way ANOVA, *p* = 0.043). Co-administration of silibinin at either 25 or 50 µM but did not reduce hepatotoxicity induced by 50 mM INH, 50 mM PZA, or a combination of INH and PZA (I/P) at 50 mM each (I/P 50/50). **(B)** Pre-administration of silibinin for 24 h, followed by the co-administration of silibinin with INH or PZA for a further 24 h, did not prevent hepatotoxicity induced by 50 mM INH, 80 mM INH, or 50 mM PZA. **(C)** Administration of INH or PZA for 24 h, followed by the administration of silibinin alone (with washout) or silibinin with INH or PZA (without washout) for a further 24 h, did not aid in the recovery of LO2 from 50 mM INH, 80 mM INH, or 50 mM PZA. Data represent mean ± S.E.M. of at least two replicates. * *p* < 0.05 vs respective vehicle controls.

### 3.2 Silibinin reduced oxidative damage of INH and PZA on classical intracellular targets

After establishing silibinin’s role as a rescue adjuvant in INH-induced hepatotoxicity, we characterised silibinin’s ability to reduce intracellular ROS levels and oxidative damage to proteins, lipids, and DNA. We assessed these intracellular indicators of oxidative stress for two reasons: they play critical roles in cellular function and survival, and their measurements have been widely studied and are well-established (Halliwell and Whiteman, 2004). These experiments showed that 50 **μ**M silibinin mitigated the increase in intracellular ROS levels when co-administered with I/P 40/10 over 24 h (**Fig. 3A**). To assess whether the attenuation of intracellular ROS production translated into a reduction in damage to important biomolecules, we then quantified the corresponding oxidative damage incurred on proteins, lipids, and DNA. These experiments revealed that 25 and 50 **μ**M silibinin significantly reduced protein carbonylation and lipid peroxidation levels (**Fig. 3B-C**). Importantly, silibinin’s reduction of oxidative stress was independent of DNA oxidative damage induced by I/P 40/10 (**Fig. 3D**) as measured using the Comet assay, which is especially useful for detecting genotoxicity because it paints a holistic picture of overall DNA damage by accounting for multiple genotoxic mechanisms (Collins, 2014). Overall, silibinin’s reduction of ROS levels led to a reduction in protein carbonylation and lipid peroxidation, but not in DNA fragmentation.

**Figure 3.**
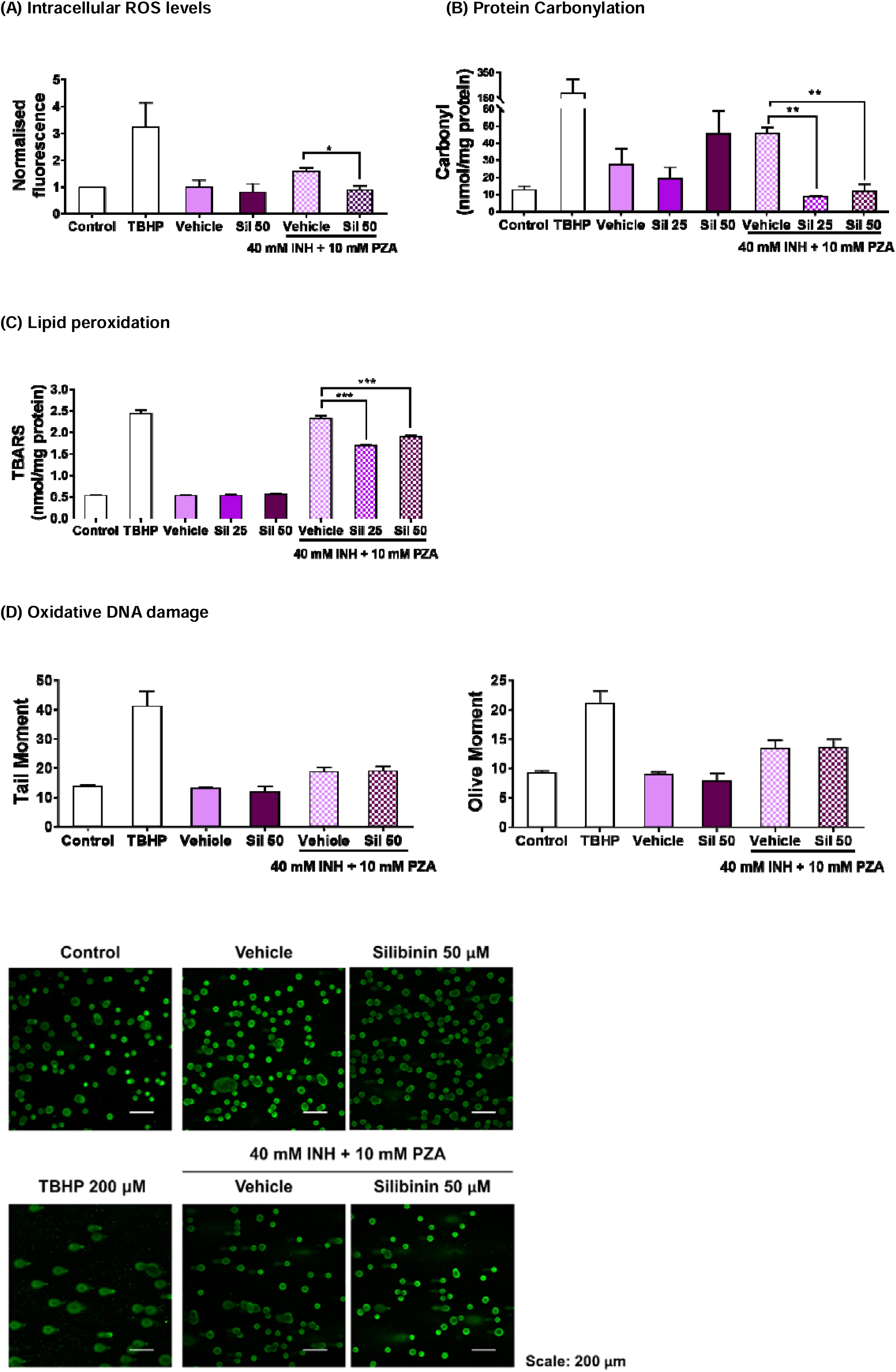
(color, 2-column). Silibinin reduced ROS levels and oxidative damage when co-administered with a combination of INH and PZA. Positive controls were treated with the oxidising agent TBHP 200 μM for 2 h. To avoid excessive hepatocyte death, the concentrations of INH and PZA were limited to 40 mM and 10 mM respectively when treated in combination (I/P 40/10) over 24 h. **(A)** 50 μM silibinin reduced intracellular ROS levels (one-way ANOVA, *p* = 0.004). **(B)** Silibinin decreased carbonylation levels, a marker of oxidative damage in proteins, at 25 μM (one-way ANOVA, *p* = 0.0011) and 50 μM (one-way ANOVA, *p* = 0.0017). **(C)** Silibinin reduced lipid peroxidation levels as measured by the TBARS assay at 25 μM (one-way ANOVA, *p* = 0.0001) and 50 μM (one-way ANOVA, *p* = 0.0010). **(D)** Silibinin’s reduction of ROS levels at 50 μM was independent of DNA oxidative damage reduction as visually assessed, and as measured quantitatively by Tail Moment and Olive Moments. Administration of silibinin alone did not trigger DNA fragmentation. Data represent mean ± S.E.M. of at least two replicates. * *p* < 0.05, ** *p* < 0.01, *** *p* < 0.001 vs vehicle control co-administered with I/P 40/10.

### 3.3 Silibinin protected against apoptosis by maintaining mitochondrial membrane potential

As oxidative stress has been reported to trigger apoptosis via caspase-9 and, subsequently, caspase-3 activation in the intrinsic pathway (Redza-Dutordoir and Averill-Bates, 2016), the drug-induced ROS levels would likely result in an increase in cell death as well. Therefore, we also measured silibinin’s ability to reduce apoptotic activity in LO2 to further reinforce the association between the observed decrease in cell viability and the increase in ROS levels. The use of caspase-3 activity to gauge apoptosis yielded two key observations (**Fig. 4A**). First, the co-administration of silibinin significantly mitigated the induction of caspase-3 activity by I/P 40/10. Second, silibinin alone did not induce caspase-3 activity. This suggests that the observed decrease in cell viability is solely attributed to the exposure of LO2 to I/P 40/10.

**Figure 4.**
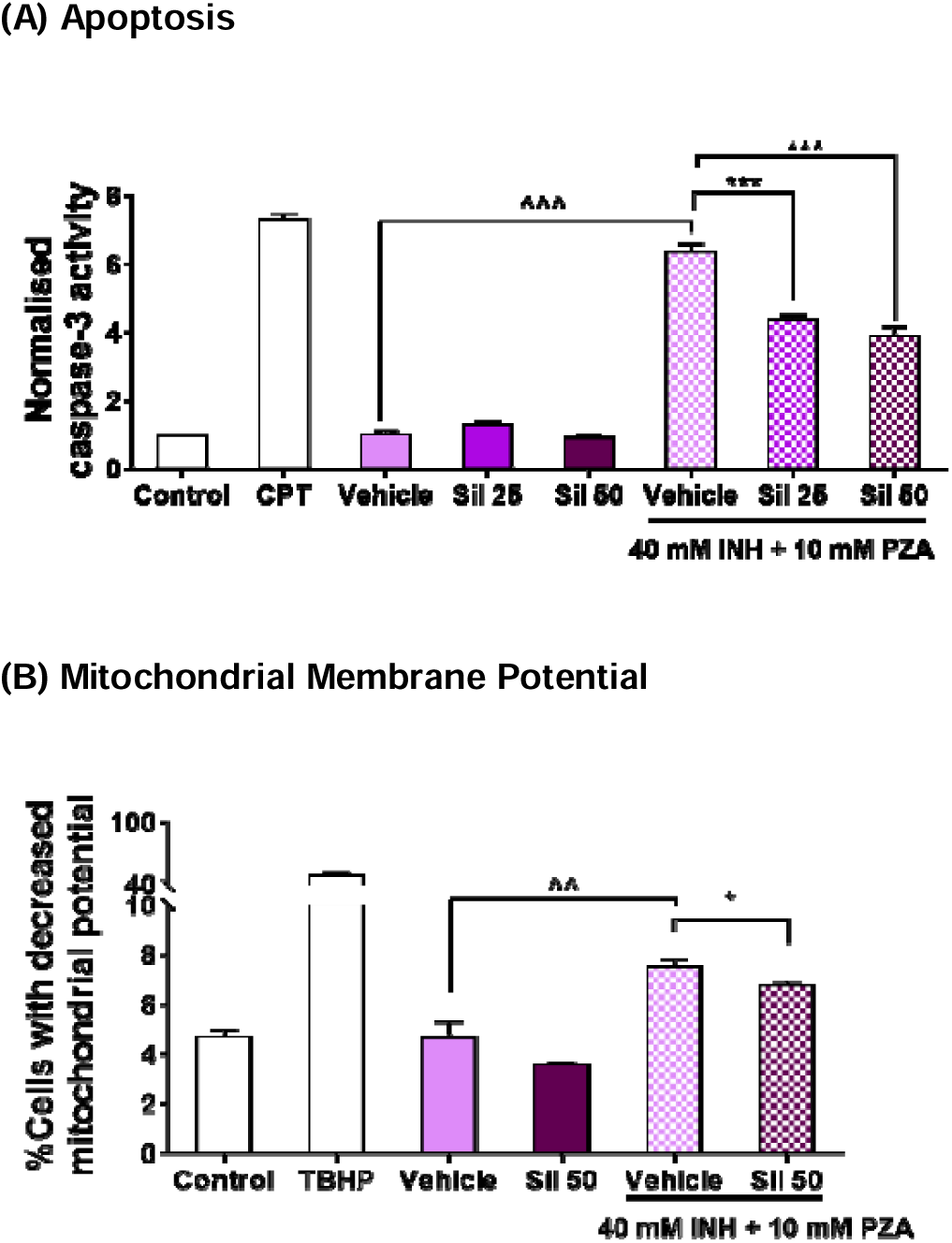
(color, 2-column). Silibinin reduced apoptosis when co-administered with a combination of INH and PZA by maintaining mitochondrial membrane potential. Various concentrations of silibinin were co-administered with I/P 40/10 over 18 h. **(A)** The administration of I/P 40/10 significantly increased caspase-3 activity (one-way ANOVA, *p* < 0.0001). The co-administration of silibinin with I/P 40/10 reduced the activity of caspase-3, the final mediator of both the intrinsic and extrinsic apoptotic pathways, when silibinin was administered at 25 μM (one-way ANOVA, *p* < 0.0001) and 50 μM (one-way ANOVA, *p* < 0.0001). The positive control was treated with camptothecin (CPT) 5 μM for 24 h. **(B)** The administration of I/P 40/10 negatively affected LO2 cells’ membrane potential (unpaired t-test, *p* = 0.0074). 50 μM silibinin reduced the percentage of cells whose membrane potential was negatively affected by I/P 40/10 (unpaired t-test, *p* = 0.0408). Positive control was treated with the oxidising agent TBHP 200 μM for 1 h. Data represent mean ± S.E.M. of three replicates. * *p* < 0.05, *** *p* < 0.001 vs vehicle control co-administered with I/P 40/10, ^^ *p* < 0.01, ^^^ *p* < 0.001 vs respective vehicle controls.

Oxidative mitochondrial stress has been identified as the key driver of INH-induced apoptotic activity: the increase in ROS levels promotes megamitochondria formation, consequently triggering cytochrome c release and upregulating apoptotic signalling (Ahadpour et al., 2016; Boelsterli and Lee, 2014). Thus, having observed that silibinin attenuated apoptotic activity, we then interrogated silibinin’s effect on preserving mitochondrial function. We observed that silibinin indeed slightly attenuated the proportion of cells with mitochondrial membrane potential transition induced by I/P 40/10 (**Fig. 4B**). This may suggest that silibinin’s reduction of oxidative stress may ameliorate mitochondrial dysfunction and, in turn, apoptotic activity.

### 3.4 Silibinin restored HO-1 expression and induced Srxn1 expression in TAMH

Another aspect of INH’s hepatotoxic effect arises from its suppression of proteins expressed in the Nrf2-ARE pathway (Chen et al., 2013; Peng et al., 2016). However, this effect has not been reported for PZA. To evaluate whether silibinin’s hepatoprotective effect entails the induction of Nrf2-ARE-related protein expression, we first profiled the individual effects of INH and PZA in LO2. In these preliminary tests, INH, but not PZA, suppressed the expression of HO-1. However, other ARE responsive genes, such as Gclc, NQO1, and Srxn1, were not suppressed by INH (**Fig. S3**). Silibinin was then co-administered with I/P 40/10 to test our hypothesis that silibinin’s utility in the PIER regimen arises from its induction of these antioxidant enzymes. Because the expressions of these four antioxidant proteins may differ across cell lines, we tested our hypothesis in both LO2 and TAMH.

While silibinin has been reported to exhibit an indirect antioxidative effect by upregulating the Nrf2-ARE pathway (Kim et al., 2012), we found that silibinin’s hepatoprotection in LO2 was independent of Gclc, HO-1, NQO1, and Srxn1 induction (**Fig. 5A**). Since the Nrf2-ARE pathway in LO2 may not be sensitive to suppression by I/P compared to other cell lines, we further verified our observation in TAMH, in which the administration of silibinin alone induced Srxn1 expression. Moreover, when co-administered with 40 mM INH, silibinin restored HO-1 expression to normal levels, though these effects were independent of Gclc and NQO1 induction (**Fig. 5B**).

**Figure 5.**
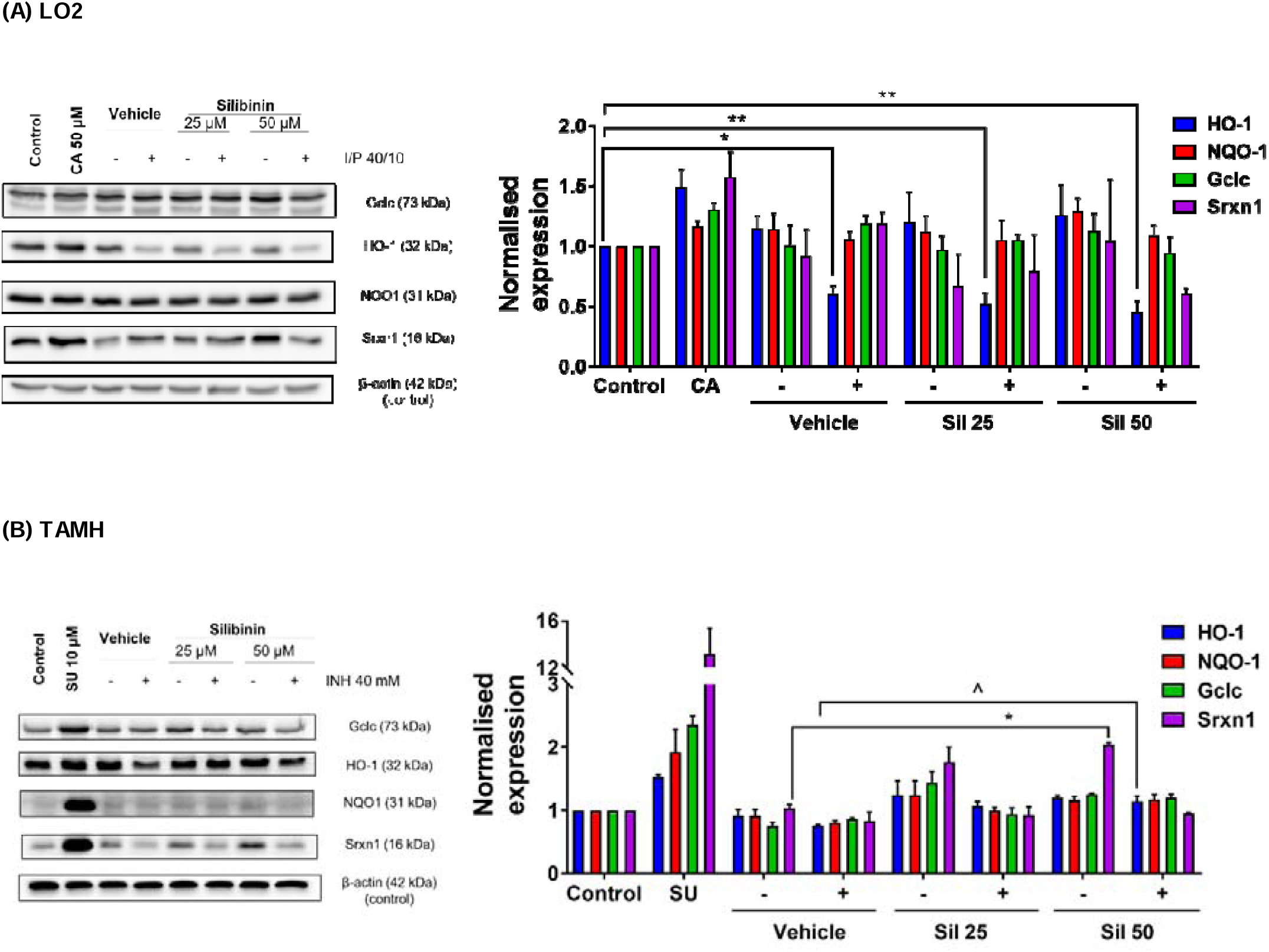
(color, 2-column). Silibinin induced expression of proteins in the Nrf2-ARE pathway and restored protein expression. Vehicle control was treated with 0.05% v/v DMSO. **(A)** In LO2, silibinin’s reduction of ROS levels when co-administered with I/P 40/10 was independent of HO-1 protein restoration. The administration of I/P 40/10 significantly reduced HO-1 levels without silibinin (one-way ANOVA, *p* = 0.0150), or with silibinin at 25 μM (one-way ANOVA, *p* = 0.0051) and 50 μM (one-way ANOVA, *p* = 0.0022). Silibinin alone did not induce the expression of Gclc, HO-1, NQO1, and Srxn1. Positive controls were treated with the Nrf2-ARE inducer CA 50 μM for 24 h. Data represent mean ± S.E.M. of three replicates. * *p* < 0.05, ** *p* < 0.01 vs negative control. **(B)** In TAMH, 50 μM silibinin alone induced Srxn1 expression (one-way ANOVA, *p* = 0.0237), but the co-administration of silibinin with 40 mM INH did not restore Srxn1 expression to pre-suppression levels. In contrast, 50 μM silibinin restored HO-1 expression (one-way ANOVA, *p* = 0.0333), but did not induce HO-1 when was administered alone. This effect did not extend to Gclc and NQO1 restoration. Positive controls were treated with the Nrf2-ARE inducer SU 10 μM for 24 h. Data represent mean ± S.E.M. of two replicates. * *p* < 0.05 vs vehicle control, ^ *p* < 0.05 vs vehicle control co-administered with hepatotoxic regimen.

## 4. Discussion

DILI is the most frequently cited reason for the withdrawal of drugs, especially when the manifestations of the hepatotoxicity are complex and require a better understanding of the underlying mechanisms (Björnsson, 2016). Therefore, we have simulated silibinin’s clinical roles in prophylaxis, rescue, and recovery of PIER-induced hepatotoxicity using an *in vitro* model with the respective pre-, co-, and post-administration of silibinin with the hepatotoxic regimens. As a prophylactic agent, silibinin would be taken before starting the PIER regimen to protect patients from future hepatotoxicity; as a rescue agent, silibinin would be co-administered with the PIER regimen to mitigate hepatotoxicity; and as a recovery agent, silibinin would be prescribed after the onset of PIER-induced hepatotoxicity to aid in the healing process. We found that silibinin was mainly useful as a rescue adjuvant (**Fig. 2A**) and ascertained that silibinin’s hepatoprotective effect arises from two aspects. First, silibinin reduces intracellular levels of oxidative stress and oxidative damage to intracellular targets and mitochondria, leading to decreased apoptotic activity (**Fig. 3A-D, Fig. 4A-B**). This observation is consistent with silibinin’s ability to reduce *in vivo* markers of direct oxidative damage, such as DNA fragmentation levels, lipid peroxidation, and mitochondrial dysfunction also reported by other authors (Patel et al., 2010; Surai, 2015; Vargas-Mendoza et al., 2014). Second, silibinin induces Nrf2-ARE-related protein expression (**Fig. 5B**). This also coincides with silibinin’s ability to increase levels of endogenous proteins that protect cells from oxidative damage, including various mediators along the MAPK pathway (Chen et al., 2006), thioredoxin (Rodriguez-Garcia et al., 2017), and superoxide dismutase (SOD) (Hamza and Al-Harbi, 2015).

When used as a rescue adjuvant, silibinin was the most significantly hepatoprotective within INH’s “Goldilocks zone” (i.e. synonymous to a zone that is neither too high or too low) (**Fig. 2A**). Specifically, the “Goldilocks zone” is a range of toxicant concentrations around its IC_50_, the toxicant concentrations that trigger DILI to approximately 50% cell viability: the IC_50_ of PZA and INH are 60 and 73 mM respectively (**Fig. S2A-B**). This implies that silibinin may be the most efficacious within a specific window of INH concentrations that are neither too severe nor too mild. This observation is consistent with other *in vitro* findings, which involve the characterisation of silibinin’s hepatoprotective effects at toxicant concentrations around the IC_50_ values of their respective assays (Mann et al., 2017; Singh et al., 2012). The existence of INH’s “Goldilocks zone” has two clinical implications in discussing silibinin’s role as a rescue adjuvant. First, it suggests that silibinin may be the most efficacious at moderate levels of DILI, and correspondingly less hepatoprotective in very early or late stages of DILI. Thus, depending on a patient’s liver function, silibinin’s dose can be carefully titrated to optimise the magnitude of hepatoprotection while reducing silibinin’s potential side effects. The optimal silibinin concentration from our viability experiments was determined to be 25 **μ**M when DILI was induced with 80 mM INH (**Fig. 2A**), which may be explained by silibinin’s pro-oxidative and pro-apoptotic effects at higher concentrations, an observation that corroborates experiments conducted by other research groups in rats (Malekinejad et al., 2012) and other *in vitro* cell lines (Mann et al., 2017; Procházková et al., 2011; Surai, 2015). In other words, increasing silibinin’s concentration may not always lead to increased hepatoprotection.

At the same time, silibinin was not useful as an adjuvant in prophylaxis and recovery (**Fig. 2B-C**), suggesting that silibinin may not prevent, or help patients recover from, PIER-induced hepatotoxicity respectively. The lack of prophylactic effect when silibinin is administered before the toxicant regimen may be attributed to the inadequate induction of antioxidant responses, especially that belonging to the Nrf2-ARE pathway (**Fig. 5A**). Our observation that silibinin did not promote the recovery process may also indicate that silibinin may not be involved in the regenerative mechanisms that restore normal hepatocyte function after DILI has occurred (Forbes and Newsome, 2016). However, silibinin has been shown to reduce stellate cell migration, which may reduce fibrotic activity (Ezhilarasan et al., 2016; Kim et al., 2012). Therefore, future directions to better characterise silibinin’s hepatoprotection in the case of recovery therapy may centre around the use of co-cultures, which can mimic paracrine responses.

Taken together, these insights generated from our viability studies may be particularly useful in an era of personalised medicine, in which clinicians can tailor bespoke regimens to patient’s conditions rapidly and accurately, instead of using a one-size-fit-all approach (Andrade, 2015).

Silibinin also protected against apoptosis induced by I/P (**Fig. 4A**) independently of viability restoration (**Fig. 2A**) by reducing intracellular oxidative stress. This reduction in I/P-induced oxidative stress manifested in two ways: decreased oxidative damage to classical intracellular targets, as measured by protein carbonylation and lipid peroxidation (**Fig. 3B-C**), and in the restoration of mitochondrial membrane potential (**Fig. 4B**). Interestingly, though silibinin has been reported to protect against doxorubicin-induced DNA oxidative fragmentation (Patel et al., 2010), silibinin did not appear to protect against oxidative DNA damage induced by I/P 40/10 in our study (**Fig. 3D**). Our observations on these four hallmarks of oxidative damage corroborate existing *in vitro* and *in vivo* studies on silibinin’s antioxidant effect in showing that silibinin mitigates the elevated lipid peroxidation and protein carbonylation levels in DILI (Song et al., 2006), and further imply that silibinin may not protect against all forms of INH- and PZA-induced oxidative damage. At the same time, our Comet assay results lend credence to silibinin’s safety at the concentrations used in our study. Thus, our observation that there was no difference in the magnitudes of the Tail Moment and Olive Moment between the control and silibinin-treated samples (**Fig. 3D**) suggests that silibinin did not induce DNA damage in our study. Coupled with the observation that silibinin did not induce caspase-3 activity (**Fig. 4A**), this reinforces silibinin’s safety profile and supports its development in further studies.

Other than functioning as a direct antioxidant, silibinin and its analogues have also been reported to induce the levels and activities of various endogenous antioxidants (Kim et al., 2012; Roubalová et al., 2017). Therefore, we chose to investigate Nrf2-ARE, a major antioxidant pathway. Silibinin’s hepatoprotective effect in LO2 was independent of Nrf2-ARE pathway activation or restoration after suppression by INH (**Fig. 5A**), agreeing with our earlier observation that silibinin did not function as a preventive agent *in vitro* (**Fig. 2B**): if silibinin had induced the protective Nrf2-ARE pathway, the upregulation of antioxidant enzymatic systems would serve to protect against INH- or PZA-induced hepatotoxicity. In contrast, silibinin induced Srxn-1 expression and restored HO-1 expression in TAMH (**Fig. 5B**). Interestingly, though we found that this was independent of silibinin’s hepatoprotection in TAMH (**Fig. S4**), our findings corroborate evidence in rats (Kim et al., 2012) and mice (Liu et al., 2019) that silibinin’s induction of Nrf2-ARE pathway may contribute towards its hepatoprotective effect in rodents.

The differences in observations made in LO2 and TAMH can be ascribed to possible Nrf2-independent mechanisms of hepatoprotection. Indeed, apart from exhibiting a direct hepatoprotective effect, silibinin may modulate pathways other than Nrf2-related upregulation of antioxidant enzymes. In fact, the hepatoprotective effect of tert-Butylhydroquinone has been ascribed to its effects on autophagy (Li et al., 2014), and a similar effect may exist in LO2. In contrast, the Nrf2-independent modulation of HO-1 expression has also been reported in cases of muscular atrophy (Kang et al., 2014). Taken together, these observations suggest that ARE expression may also be controlled by less understood constitutive pathways besides the well-established induction by Nrf2 activation (Li et al., 2019).

An alternative explanation for this interesting phenomenon is that the Nrf2-ARE pathway is activated by silibinin’s metabolites, rather than silibinin itself. As TAMH is metabolically active (Davis and Stamper, 2016), it may convert silibinin to metabolites that structurally resemble the analogue 2,3-dehydrosilydianin, which has been reported to upregulate NQO1 activity (Roubalová et al., 2017). The differences between our results and previous findings (Kim et al., 2012; Roubalová et al., 2017) may therefore be attributed to innate metabolic, transporter-related, and physiological differences between various cell lines. Our choice of LO2 has its distinct advantages: Not only is LO2 more representative of human liver physiology than HepG2 (Wu et al., 2016), but LO2 also expresses higher levels of CYP2E1 than HepaRG that enable LO2 to convert INH to its toxic metabolite hydrazine (Liang et al., 2010). Notably, HepaRG’s poor expression of CYP2E1 has cast doubt on its relevance in INH-induced hepatotoxicity models (Du et al., 2016). Concurrently, our *in vitro* experiments using human cell lines serve as useful cross-references for other *in vitro* (Mann et al., 2017; Singh et al., 2012), *in vivo* animal (Srivastava et al., 2008), and human (Luangchosiri et al., 2015) studies on silibinin’s protection against PIER-induced hepatotoxicity.

Our work on human-relevant LO2 thus buttresses these reports on silibinin’s hepatoprotective ability by showing that silibinin reduces hepatotoxicity induced by INH, and further clarifies the mechanisms of PZA- and INH-induced hepatotoxicity. The observation that INH suppressed HO-1 expression in LO2 when used in combination with PZA (**Fig. 5A**) is consistent with the current paradigm in which INH reduces both the mRNA transcription levels and the activity of HO-1 (Chen et al., 2013; Peng et al., 2016). By suppressing HO-1 expression, INH increases LO2 cells’ susceptibility to oxidative stress mediated by the increase in intracellular ROS levels (**Fig. 3A**). In contrast, because INH suppressed Gclc, NQO1, and Srxn1 expression in TAMH but not in LO2, the observed decrease in viability in LO2 (**Fig. 2A**) is likely independent of the expression levels of these three proteins. This also suggests that Gclc, NQO1, and Srxn1 may play a smaller role in mitigating oxidative stress induced in LO2 by I/P.

At the same time, we observed that silibinin was more hepatoprotective against INH than PZA or I/P. This observation appears to suggest that silibinin may not mitigate mechanisms involved in PZA-induced hepatotoxicity *in vitro*. Therefore, silibinin may need to be used carefully as a rescue adjuvant in the overall PIER regimen, which combines the use of INH and PZA. In fact, silibinin may be more useful in triple ATT regimens that exclude the use of PZA, which are often used in patients who suffer from hepatotoxicity (Wang et al., 2016b). Indeed, PZA-induced hepatotoxicity is complex and poorly understood. Despite PZA’s greater association with hepatotoxicity than INH, research on hepatotoxicity has mostly centred on the latter, and the mechanisms responsible for INH-induced hepatotoxicity are becoming more well understood in recent years (Ramachandran et al., 2018). Specifically, oxidative stress arising from the toxic INH metabolite hydrazine has been validated as a major mechanism in INH-induced hepatotoxicity using pharmacodynamic and pharmacokinetic evidence (Boelsterli and Lee, 2014; Mitchell et al., 1975; Ramachandran et al., 2018). In contrast, while several PZA’s metabolites have been identified (Shih et al., 2013), research characterising their toxicities has only just started emerging (Cao et al., 2018). Recently, 5-hydroxypyrazinoic acid, a metabolite of PZA, was proposed to be primarily responsible for PZA’s toxicity (Cao et al., 2018; Shih et al., 2013). However, while the conversion of PZA to 5-hydroxipyrazinoic acid is mediated by xanthine oxidase, silibinin did not protect against PZA-induced hepatotoxicity in our study (**Fig. 2A-C**) despite being a xanthine oxidase inhibitor (Surai, 2015; Varga et al., 2006). Therefore, silibinin’s inability to protect against PZA-induced hepatotoxicity may be attributed to other injury pathways that lie beyond silibinin’s hepatoprotective mode of action.

## 5. Conclusion

In summary, we have assessed and characterised silibinin’s various roles as an adjuvant in protecting against PZA- and INH-induced hepatotoxicity. Our *in vitro* experiments suggest that silibinin may be safe and efficacious as a rescue adjuvant, both fundamental considerations in the use of any drug. Further optimisation of our *in vitro* model may also enhance silibinin’s hepatoprotective effect in rescue, prophylaxis, and recovery. Using this model, we have gleaned important mechanistic insights on its hepatoprotective effect and identified novel antioxidant targets in ameliorating PIER-induced hepatotoxicity. Future directions will involve exploring the two main mechanisms by which silibinin may ameliorate hepatotoxicity; the proof-of-concept demonstrated in this project will inform subsequent *in vitro* and *in vivo* preclinical studies. Given the lack of alternative treatments in tuberculosis, the need to preserve our remaining antibiotics is paramount. These high stakes necessitate future efforts to support our preliminary work—making silibinin more clinically relevant to patients and healthcare professionals alike.

## Supporting information

Highlights

## Abbreviations

ARE: Antioxidant response element;
ATT: Antitubercular therapy;
CA: Trans-cinnamaldehyde;
CAT: Catalase;
CPT: Camptothecin;
DCFDA: 6-chloromethyl-2’,7’-dichlorodihydrofluorescein diacetate;
DILI: Drug-induced liver injury;
DMEM: Dulbecco’s minimum essential medium;
EMB: Ethambutol;
I/P: A combination of isoniazid and pyrazinamide;
INH: Isoniazid;
MTT: 3-(4,5-dimethylthiazol-2-yl)-2,5-diphenyltetrazolium bromide;
Nrf2: Nuclear factor (erythroid-derived 2)-like 2;
NQO1: NAD(P)H quinone dehydrogenase 1;
PZA: Pyrazinamide;
SOD: Superoxide Dismutase;
SU: Sulphoraphane;
TBARS: Thiobarbituric acid reactive substances;
TBHP: Tert-butyl hydroperoxide;
TCA: Trichloroacetic acid;
TMRM: Tetramethylrhodamine-methyl-ester;

## Acknowledgements

This work was supported by the National University of Singapore Grants [R148-000-217-112] and [R148-000-272-114]. The authors are grateful to A/Prof. Victor Yu for the kind gifts of various laboratory materials, A/Prof. Eng Hui Chew for the kind gifts of sulphoraphane and trans-cinnamaldehyde, A/Prof. Wai Keung Chui for the generous loan of his laboratory space and equipment, and David Packiaraj Sheela from the Drug Development Unit and Shu Ying Lee from the Confocal Microscopy Unit for imaging assistance.

This manuscript has been released as a Pre-Print at BioRxiv: Goh, Z.-H.; Tee, J. K.; Ho, H. K. bioRxiv 2019, 815241. doi:10.1101/815241.

## Statement of contribution

ZG and HH contributed conception and design of the study; ZG organized the database; ZG and HH performed the statistical analysis; ZG wrote the first draft of the manuscript; ZG, HH, and JT wrote sections of the manuscript. All authors contributed to manuscript revision, read and approved the submitted version.

---

**Figure S1.**
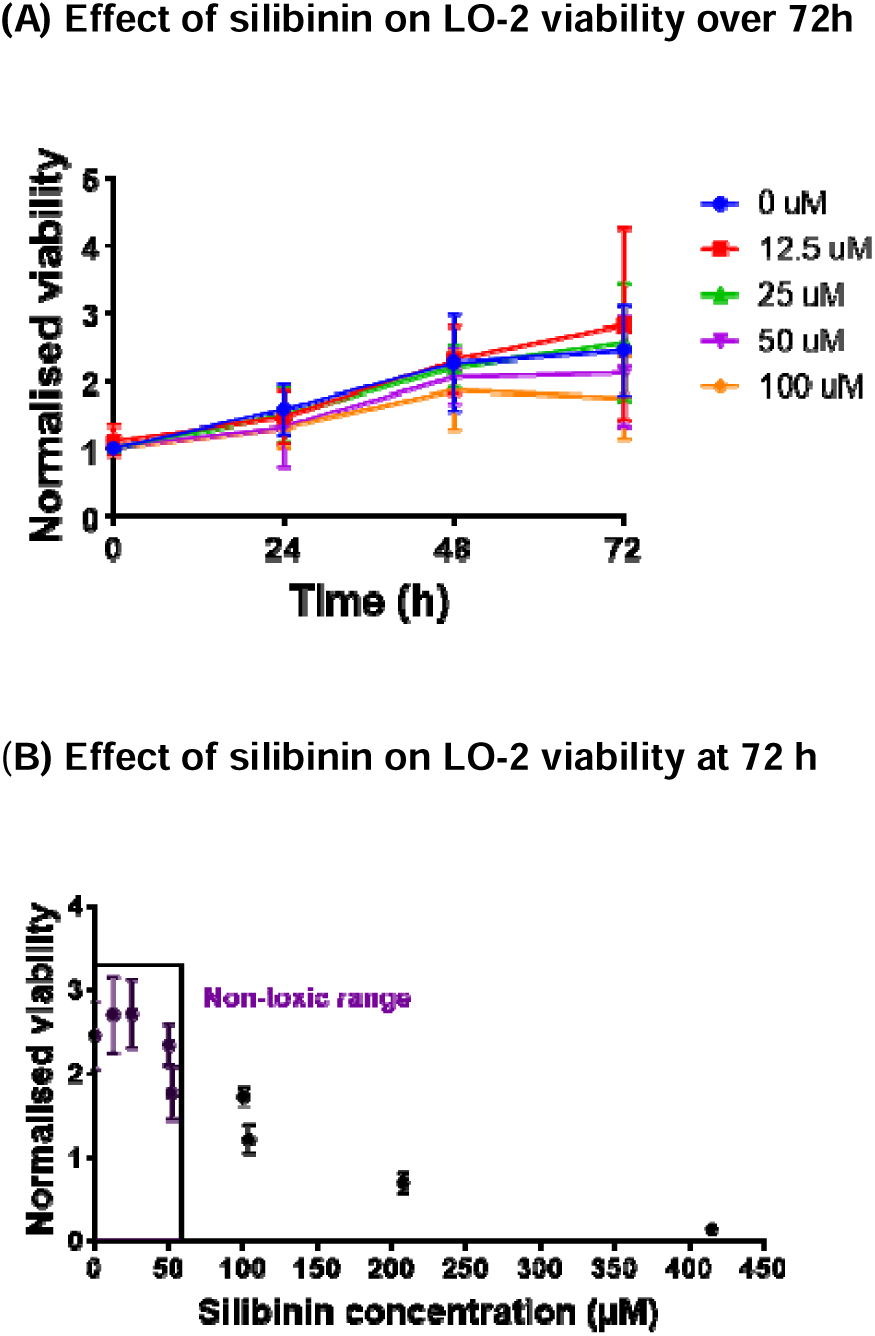
(color, 2-column). Silibinin did not affect LO2 viability at concentrations up to 50 μM over 72 h. **(A)** Silibinin at concentrations up to 50 μM over 72 h did not hinder LO2 growth. **(B)** After 72 h had passed, silibinin at concentrations up to 50 μM did not hinder LO2 growth, and silibinin above concentrations of 200 μM significantly reduced LO2 viability. Data represent mean ± S.E.M. of at least two replicates.

**Figure S2.**
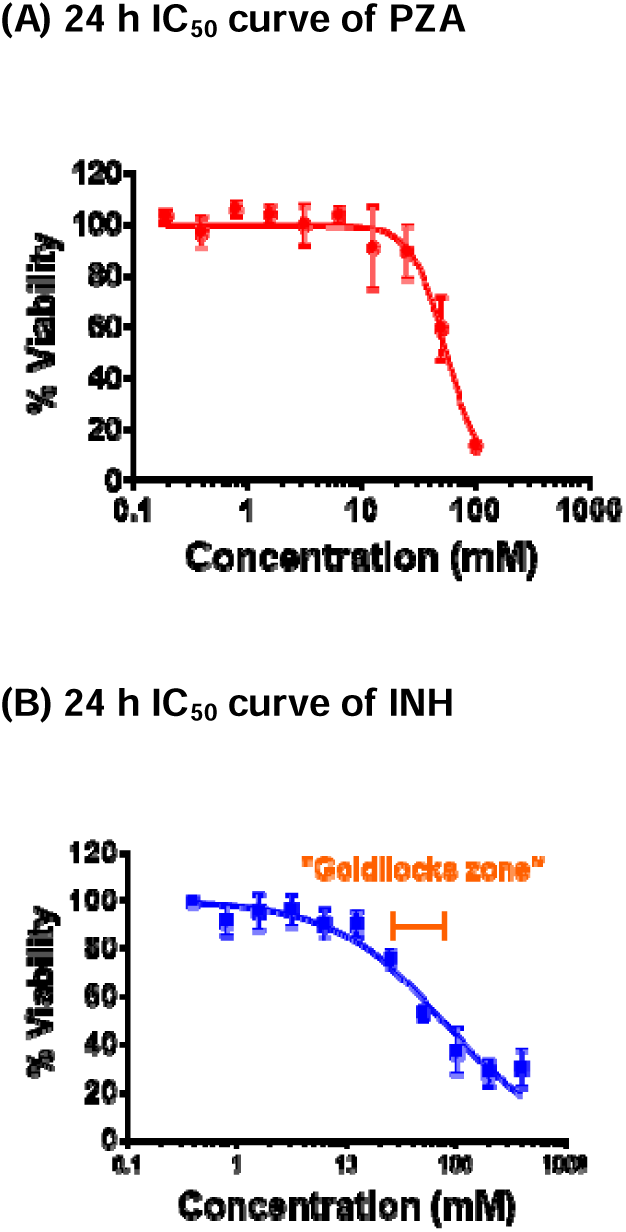
(color, 2-column). PZA and INH reduced LO2 viability when administered over 24 h. **(A)** The mean IC_50_ of PZA was 60 mM. **(B)** The mean IC_50_ of INH was 73 mM. The “Goldilocks zone” (orange) contained the concentrations of INH that induced toxicity to approximately 50% viability. The more representative IC_50_ curve of two biological replicates is shown for PZA and INH respectively. Data represent mean ± S.D. of six technical replicates.

**Figure S3.**
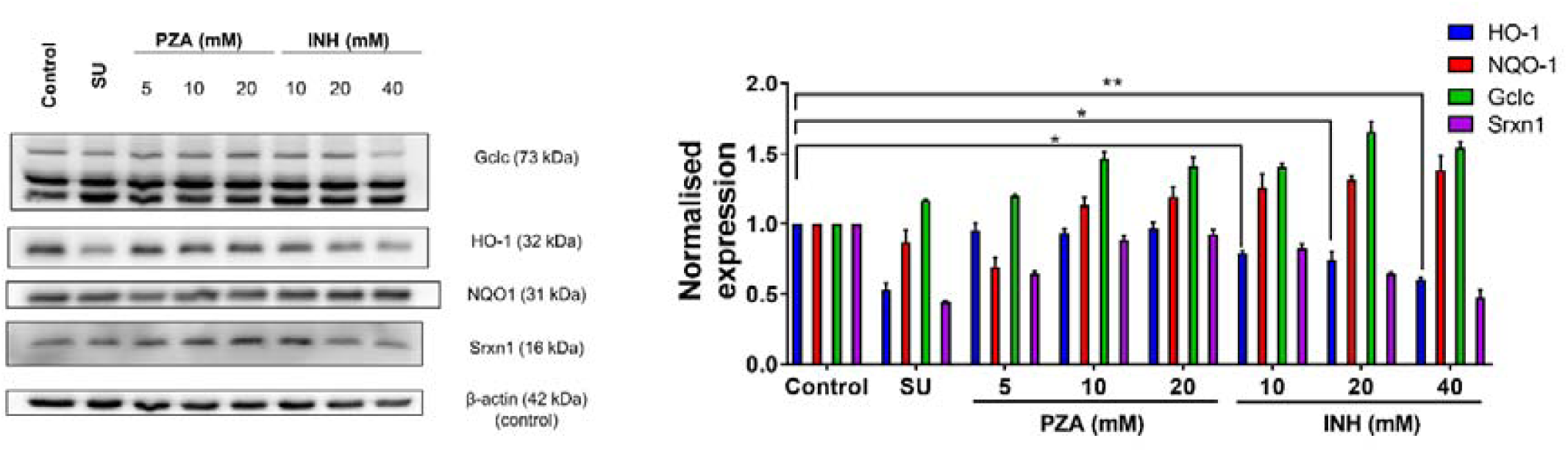
(color, 2-column). INH, but not PZA, significantly suppressed HO-1 expression in LO2. INH suppressed HO-1 expression at 10 mM (one-way ANOVA, *p* = 0.0275), 20 mM (one-way ANOVA, *p* = 0.0133), and 40 mM (one-way ANOVA, *p* = 0.0028); but not Gclc, NQO1, and Srxn1 expression in LO2 *in vitro*. PZA did not have any effect on HO-1, Gclc, NQO1, and Srxn1 at the concentrations tested. Positive control was treated with 10 μM SU for 24 h. Data represent mean ± S.E.M. of two replicates. * *p* < 0.05, ** *p* < 0.01 vs control.

**Figure S4.**
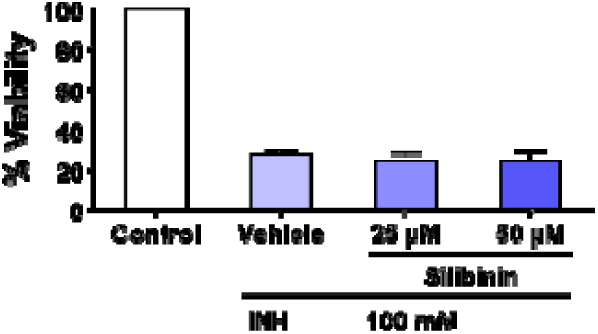
(color, 2-column). Silibinin was not useful as a rescue adjuvant in TAMH. The co-administration of silibinin at 25 and 50 μM did not preserve TAMH function when hepatotoxicity was induced using 100 mM INH. Data represent mean ± S.E.M. of three replicates.

