## Supplementary material for "An Evaluation of the *In Vitro* Roles and Mechanisms of Silibinin in Reducing Pyrazinamide- and Isoniazid-Induced Hepatocellular Damage": Highlights

- Silibinin preserved cell viability when co-administered with isoniazid
- Silibinin reduced oxidative damage induced by isoniazid and pyrazinamide
- Silibinin maintained mitochondria membrane potential, reducing apoptosis
- Silibinin activated the Nrf2-ARE pathway
