## Supplemental Figure S1 for "An Evaluation of the *In Vitro* Roles and Mechanisms of Silibinin in Reducing Pyrazinamide- and Isoniazid-Induced Hepatocellular Damage"

**(A) Effect of silibinin on LO-2 viability over 72h**

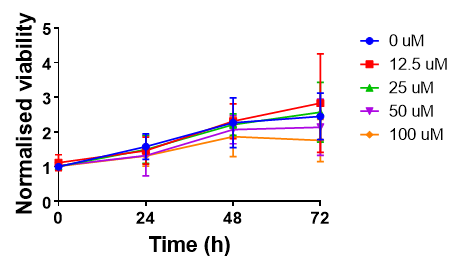

(**B) Effect of silibinin on LO-2 viability at 72 h**

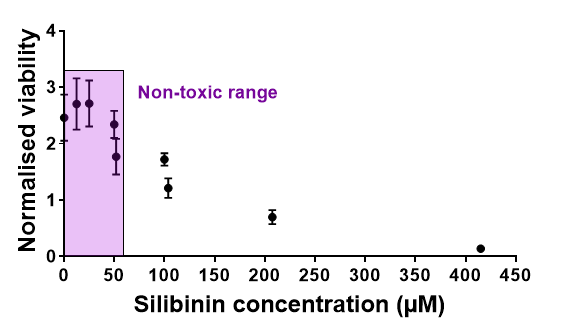

**Figure S1 (color, 2-column). Silibinin did not affect LO2 viability at concentrations up to 50 μM over 72 h. (A)** Silibinin at concentrations up to 50 μM over 72 h did not hinder LO2 growth. **(B)** After 72 h had passed, silibinin at concentrations up to 50 μM did not hinder LO2 growth, and silibinin above concentrations of 200 μM significantly reduced LO2 viability. Data represent mean ± S.E.M. of at least two replicates.
