## Supplemental Figure S2 for "An Evaluation of the *In Vitro* Roles and Mechanisms of Silibinin in Reducing Pyrazinamide- and Isoniazid-Induced Hepatocellular Damage"

**(A****) 24 h IC_50_ curve of PZA**

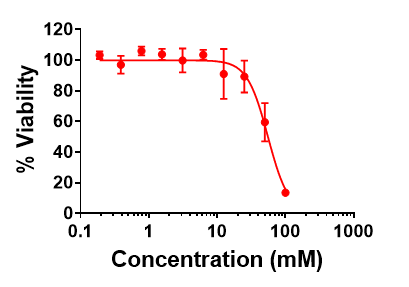

**(B) 24 h IC_50_ curve of INH**

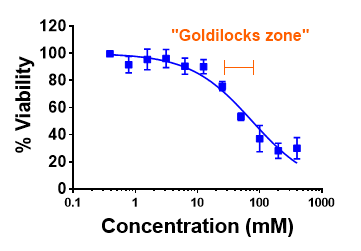

**Figure S2 (color, 2-column). PZA and INH reduced LO2 viability when administered over 24 h. (A)** The mean IC_50_ of PZA was 60 mM. **(B)** The mean IC_50_ of INH was 73 mM. The “Goldilocks zone” (orange) contained the concentrations of INH that induced toxicity to approximately 50% viability. The more representative IC_50_ curve of two biological replicates is shown for PZA and INH respectively. Data represent mean ± S.D. of six technical replicates.
