## Supplemental Figure S3 for "An Evaluation of the *In Vitro* Roles and Mechanisms of Silibinin in Reducing Pyrazinamide- and Isoniazid-Induced Hepatocellular Damage"

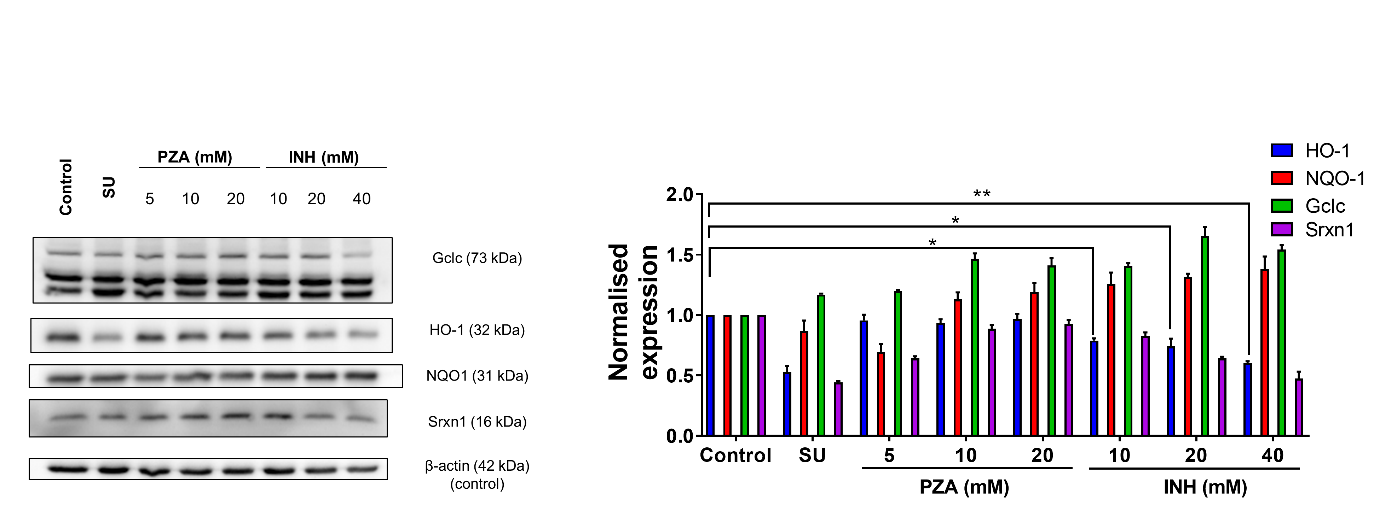

**Figure S3 (color, 2-column). INH, but not PZA, significantly suppressed HO-1 expression in LO2.** INH suppressed HO-1 expression at 10 mM (one-way ANOVA, *p* = 0.0275), 20 mM (one-way ANOVA, *p* = 0.0133), and 40 mM (one-way ANOVA, *p* = 0.0028); but not Gclc, NQO1, and Srxn1 expression in LO2 *in vitro*. PZA did not have any effect on HO-1, Gclc, NQO1, and Srxn1 at the concentrations tested. Positive control was treated with 10 μM SU for 24 h. Data represent mean ± S.E.M. of two replicates. * *p* < 0.05, ** *p* < 0.01 vs control.
