## Supplemental Figure S4 for "An Evaluation of the *In Vitro* Roles and Mechanisms of Silibinin in Reducing Pyrazinamide- and Isoniazid-Induced Hepatocellular Damage"

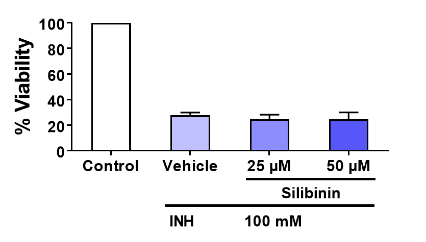

**Figure S4 (color, 2-column). Silibinin was not useful as a rescue adjuvant in TAMH.** The co-administration of silibinin at 25 and 50 μM did not preserve TAMH function when hepatotoxicity was induced using 100 mM INH. Data represent mean ± S.E.M. of three replicates.
